# Uncertainty Reduction in Biochemical Kinetic Models: Enforcing Desired Model Properties

**DOI:** 10.1101/427716

**Authors:** Ljubisa Miskovic, Jonas Béal, Michael Moret, Vassily Hatzimanikatis

## Abstract

A persistent obstacle for constructing kinetic models of metabolism is uncertainty in the kinetic properties of enzymes. Currently, available methods for building kinetic models can cope indirectly with uncertainties by integrating data from different biological levels and origins into models. In this study, we use the recently proposed computational approach iSCHRUNK (<u>i</u>n <u>S</u>ilico Approach to <u>Ch</u>aracterization and <u>R</u>eduction of <u>Un</u>certainty in the <u>K</u>inetic Models), which combines Monte Carlo parameter sampling methods and machine learning techniques, in the context of Bayesian inference. Monte Carlo parameter sampling methods allow us to exploit synergies between different data sources and generate a population of kinetic models that are consistent with the available data and physicochemical laws. The machine learning allows us to data-mine the *a priori* generated kinetic parameters together with the integrated datasets and derive posterior distributions of kinetic parameters consistent with the observed physiology. In this work, we used iSCHRUNK to address a design question: can we identify which are the kinetic parameters and what are their values that give rise to a desired metabolic behavior? Such information is important for a wide variety of studies ranging from biotechnology to medicine. To illustrate the proposed methodology, we performed Metabolic Control Analysis, computed the flux control coefficients of the xylose uptake (XTR), and identified parameters that ensure a rate improvement of XTR in a glucose-xylose co-utilizing *S. cerevisiae* strain. Our results indicate that only three kinetic parameters need to be accurately characterized to describe the studied physiology, and ultimately to design and control the desired responses of the metabolism. This framework paves the way for a new generation of methods that will systematically integrate the wealth of available omics data and efficiently extract the information necessary for metabolic engineering and synthetic biology decisions.

**Author Summary:** Kinetic models are the most promising tool for understanding the complex dynamic behavior of living cells. The primary goal of kinetic models is to capture the properties of the metabolic networks as a whole, and thus we need large-scale models for dependable *in silico* analyses of metabolism. However, uncertainty in kinetic parameters impedes the development of kinetic models, and uncertainty levels increase with the model size. Tools that will address the issues with parameter uncertainty and that will be able to reduce the uncertainty propagation through the system are therefore needed. In this work, we applied a method called iSCHRUNK that combines parameter sampling and machine learning techniques to characterize the uncertainties and uncover intricate relationships between the parameters of kinetic models and the responses of the metabolic network. The proposed method allowed us to identify a small number of parameters that determine the responses in the network regardless of the values of other parameters. As a consequence, in future studies of metabolism, it will be sufficient to explore a reduced kinetic space, and more comprehensive analyses of large-scale and genome-scale metabolic networks will be computationally tractable.

## Introduction

Kinetic models are one of the cornerstones of rational metabolic engineering as they allow us to capture the dynamic behavior of metabolism and to predict dynamic responses of living organisms to genetic and environmental changes. With reliable kinetic models, metabolic engineering and synthetic biology strategies for improvement of yield, titer, and productivity of the desired biochemical can be devised and tested *in silico* (1). The scientific community has acknowledged the utility and potential of kinetic models, and efforts towards building large- and genome-scale kinetic models were recently intensified (2–9). Nevertheless, the development of these models is still facing challenges, such as partial experimental observations and large uncertainties in available data (10–12).

The major difficulty in determining parameters of kinetic models are uncertainties associated with: (i) flux values and directionalities (13–16); (ii) metabolite concentration levels and thermodynamic properties (13–16); and (iii) kinetic properties of enzymes (2, 17). As a result of interactions of metabolite concentrations and metabolic fluxes through thermodynamics and kinetics, these uncertainties make parameter estimation difficult. Quantifying these uncertainties and determining how they propagate to the parameter space is essential for identification of parameters that should be measured or estimated to reduce the uncertainty in the output quantities such as time evolution of metabolites or control coefficients (18, 19).

In biological systems, large uncertainties and partial experimental data commonly result in a population instead of in a unique set of parameter values that could describe the experimental observations. Such population of parameter sets is typically computed using Monte Carlo sampling techniques (3–5, 8, 9, 11, 20–28). However, the problem is when certain properties differ among models in a model population. For example, one such property is flux control coefficients (FCCs)(18, 19, 29). In (30), we used the ORACLE framework (3, 4, 8, 10, 11, 31, 32) to compute a population of kinetic models along with the corresponding flux control coefficients with the aim of improving xylose uptake rate (XTR) of a glucose-xylose co-utilizing *S. cerevisiae* strain. We have found that in the same population of models that are consistent with the observed physiology FCCs can be different due to lack of data about kinetic parameters. This can lead to erroneous or conflicting conclusions and decisions about the system in metabolic engineering and synthetic biology studies.

In this contribution, to resolve such issues, we propose to formulate these problems as parameter classification: identify which of the parameters, if any, should be constrained so that the values of studied properties, such as FCCs, are in predefined ranges. For this purpose, we extended the capabilities of iSCHRUNK (<u>i</u>n <u>S</u>ilico Approach to <u>Ch</u>aracterization and <u>R</u>eduction of <u>Un</u>certainty in the <u>K</u>inetic Models), a recently introduced machine learning approach that characterizes uncertainties in parameters of kinetic models, and identifies accurate and narrow ranges of parameters that can describe a studied physiological state (17). In iSCHRUNK, machine learning is combined with methods that generate populations of kinetic models (3–5, 8, 9, 11, 20–28) to data-mine the integrated data and observed physiology together with the kinetic parameters. The extended iSCHRUNK workflow is amenable for identifying parameters that give rise to a wide variety of properties of metabolic responses. The identified parameters can further be refined in an iterative way using the stratified sampling. Moreover, a set of improvements in the parameter classification procedure was introduced to improve the classification accuracy and to allow for dealing with uncertainties in alternative physiologies, e.g., when multiple metabolite concentrations vectors are consistent with the observed physiology.

As an illustration of the capabilities of the extended iSCHRUNK, we identified the enzymes and their kinetic parameters that determine consistent FCC values related to XTR. Our results showed that by constraining only three parameters, corresponding to xylose reductase (XRI) and ATP synthase (ASN), consistent FCCs can be obtained for models computed around multiple steady-state metabolite concentrations. We further showed how the parameter classification can be improved to more accurately identify the parameter subspaces that lead to well-determined model properties.

## Results

### Uncertainty in the xylose uptake responses to genetic manipulations

In (30), we analyzed the improvement of the xylose uptake rate (XTR) during mixed glucose-xylose utilization in a recombinant *Saccharomyces cerevisiae* strain. Here, we revisited that study and built the kinetic model of *S. cerevisiae* metabolic network around the reference steady-state of metabolic fluxes and metabolite concentrations (Methods). The model contains 258 parameters and describes 102 reactions and 96 intracellular metabolites distributed over cytosol, mitochondria and extracellular environment. The experimentally determined values of kinetic parameters were missing, and the analyzed system was underdetermined, i.e., we had 102+96 computed values for steady-state fluxes and metabolite concentrations versus 258 unknown parameters. This meant that a multitude of parameter sets could reproduce the observed physiology, and we used the ORACLE framework that employs Monte Carlo sampling to generate a population of 200’000 kinetic models. We computed the flux control coefficients (FCCs) of the metabolic network and used them to rank enzymes according to their control over XTR, i.e., the highest ranked enzymes were the ones with the largest magnitude FCCs with respect to XTR. Among the top ranked enzymes, hexokinase (HXK), non-growth associated maintenance (ATPM), and NADPH reductase (NDR) had ambiguous control over XTR (Fig 1A). The distributions of the control coefficients of XTR with respect to HXK, ATPM and NDR (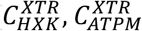, and 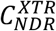, respectively) were extensively spread around zero, and we could not deduce with certainty whether the control of these enzymes over XTR was positive or negative.

**Fig 1.**
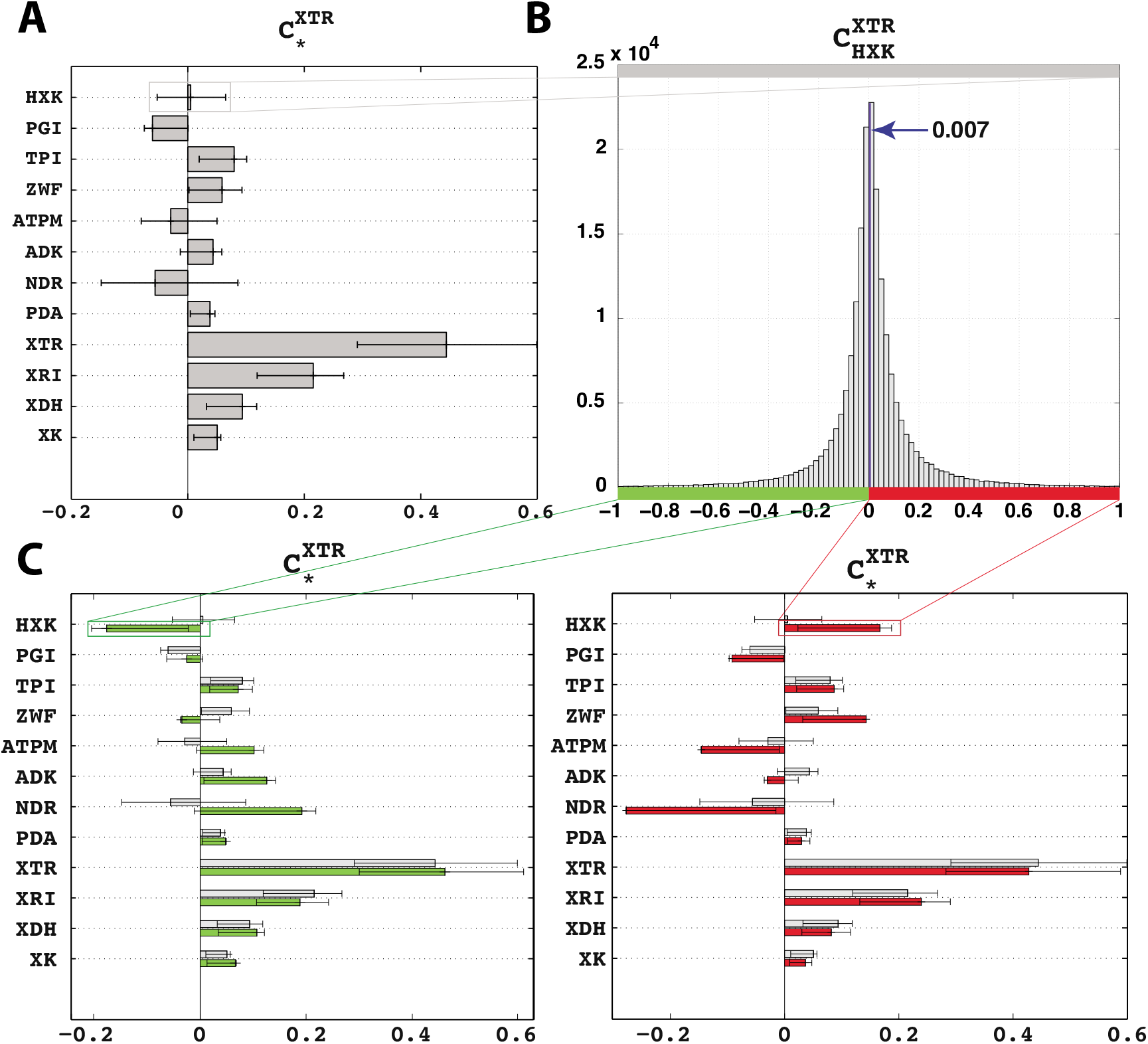
Ambiguous control of HXK, ATPM and NDR over xylose uptake (XTR) during mixed glucose-xylose fermentation. **(A)** Control coefficients of the top enzymes over XTR. The bars represent the mean values of the control coefficients through XTR. The error bars denote the 1^st^ and the 3^rd^ quartile of the control coefficients with respect to their mean value, i.e., 50% of the samples closest to the mean value are between the error bars. **(B)** The distribution of the control coefficient of HXK over XTR was centered around zero. **(C)** Pruned population of the control coefficients containing only models that had a negative control of HXK over XTR (left, green bars) or a positive one (right, red bars). For comparison purposes, the non-pruned population of control coefficients is also shown (left and right, gray bars). Enzymes: *HXK*, hexokinase; *PGI*, glucose-6-phosphate isomerase; *TPI*, triose phosphate isomerase; *ZWF*, glucose-6-phosphate-1-dehydrogenase; *ATPm*, non-growth associated ATP maintenance; *ADK*, adenylates kinase; *NDR*, NADPH reductase; *PDA*, pyruvate dehydrogenase; *XTR*, xylose transporters; *XRI*, xylose reductase; *XDH*, xylitol reductase; *XK*, xylulokinase; The complete list of enzymes and chemical species is provided in S1 File.

The population of control coefficients 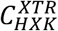 was nearly symmetric around zero with a mean of 0.005 and 47% of samples had negative values (Fig 1B). We split the population of kinetic models based on the sign of 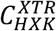, and we analyzed the two populations with a negative (Fig 1C, left) and a positive (Fig 1C, right) control of HXK over XTR. The split in the population did not have a substantial effect on the majority of the control coefficients. Interestingly, the exceptions were precisely the other enzymes with the ambiguous control over XTR, i.e., ATPM and NDR, which exhibited a negative correlation with HXK (Fig 1C). This suggested that there were two distinct populations of kinetic models. The fact that models within these two populations have several common metabolic responses further implied that each of these two populations of models had distinct values of some kinetic parameters that determined such metabolic responses.

### Identification of significant parameters determining control of HXK over XTR

We used the Classification and Regression Trees (CART) algorithm (33, 34) to identify significant parameters that determine responses of XTR to changes in HXK activity. The CART algorithm partitions the parameter space into hyper-rectangles determined by the ranges of parameters that satisfy the studied property. Here, we used as parameters the degree of saturation of the enzyme active site, σ_A_ (10), because this quantity is constrained in a well-defined range between 0 and 1 (Methods), and the desired property was the negative control of HXK over XTR. The inputs of parameter classification were: (i) the information for each out of 200’000 parameter sets whether or not it gave rise to the negative control of HXK over XTR; and (ii) parameter values of 200’000 parameter sets. Subsequently, we will refer to hyper-rectangles computed by the CART algorithm as *rules*.

To measure the performance of parameter classification we defined the *performance index* (PI), which quantifies a portion of parameter sets giving rise to the studied property. In this work, out of all parameter sets that satisfy rules (or a rule) inferred by parameter classification, PI quantifies how many of them are giving rise to the negative control of HXK over XTR. For example, within a population of models satisfying a rule, if 40% of models give rise to the negative control of HXK over XTR, then PI of this rule is 0.4.

#### Reduced number of parameters determine control of HXK over XTR

We performed parameter classification on 200’000 parameter sets of 258 parameters, and the algorithm identified 76 rules. In the identified rules, only 46 out of 258 parameters were constrained, whereas the remaining parameters had no effect on the control of HXK over XTR, i.e., their σ_A_ values could take any value between 0 and 1 (Methods). Kinetic subspaces defined by these rules had a portion of parameter sets giving rise to the negative control ranging from PI=0.50 to 0.78, i.e., 50-78% of parameter sets satisfying these rules resulted in the negative control (S2 File). This was a noteworthy improvement compared to the overall kinetic space with 47% of such parameter sets.

#### Preselection and identification of significant parameters

Our finding that a reduced number of parameters determines control of HXK over XTR suggested that statistical methods, such as Fisher’s linear discriminant score (35, 36), can be used to preselect the significant parameters, i.e., the parameters that affect the studied property. Fisher’s linear discriminant score allows us to quickly preselect parameters by analyzing the parameter distributions (Methods). We preselected 79 (out of 258) parameters that passed the threshold of 1% of the maximal Fisher’s linear discriminant score (Methods), and the values of these 79 parameters in 200’000 parameter sets were then used in the parameter classification. The classification algorithm inferred 78 rules, and remarkably, 70 of these rules coincided with the ones obtained in the first study (S3 Fig and S2 File). As expected, the ranges of the obtained PIs also coincided in the two studies. This result indicated that Fisher’s linear discriminant score is a good measure for identifying significant parameters and we used this score for parameter preselection in all further studies.

The 78 rules obtained from the study with the preselected parameters were defined by constraints on 39 parameters that corresponded to only 24 enzymes (S2 File). No rule was defined with more than 13 parameters and less than four parameters. We ranked the rules in the descending order according to the number of parameter sets that satisfy them (Methods). The top rule was defined by constraints on eight parameters, and it enclosed a subspace with 9285 parameter sets and PI of 0.65 (Table 1). The 2^nd^ and 3^rd^ ranked rules had higher values of PI than the 1^st^ ranked rule (0.73 for the 2^nd^ and 0.76 for the 3^rd^ rule), but smaller subspaces were enclosed (8049 and 6342 parameter sets for the 2^nd^ and the 3^rd^ rule, respectively). As expected, the distributions of control coefficients of XTR with respect to HXK corresponding to the parameter sets that satisfied Rules 1, 2 and 3 were biased toward negative values (Fig 2). Indeed, compared to the distribution for the overall population of parameters with the mean of 0.005 and the median of 0.005, the distributions corresponding to the three rules were shifted toward the negative values with the means of −0.082, −0.111 and −0.132, and the medians of −0.024, −0.036 and −0.044 for Rule 1, Rule 2 and Rule 3, respectively (Fig 2). These results demonstrate that the parameter classification algorithm can reliably be used to identify the significant parameters and their ranges that give rise to the negative control of HXK over XTR.

**Fig 2.**
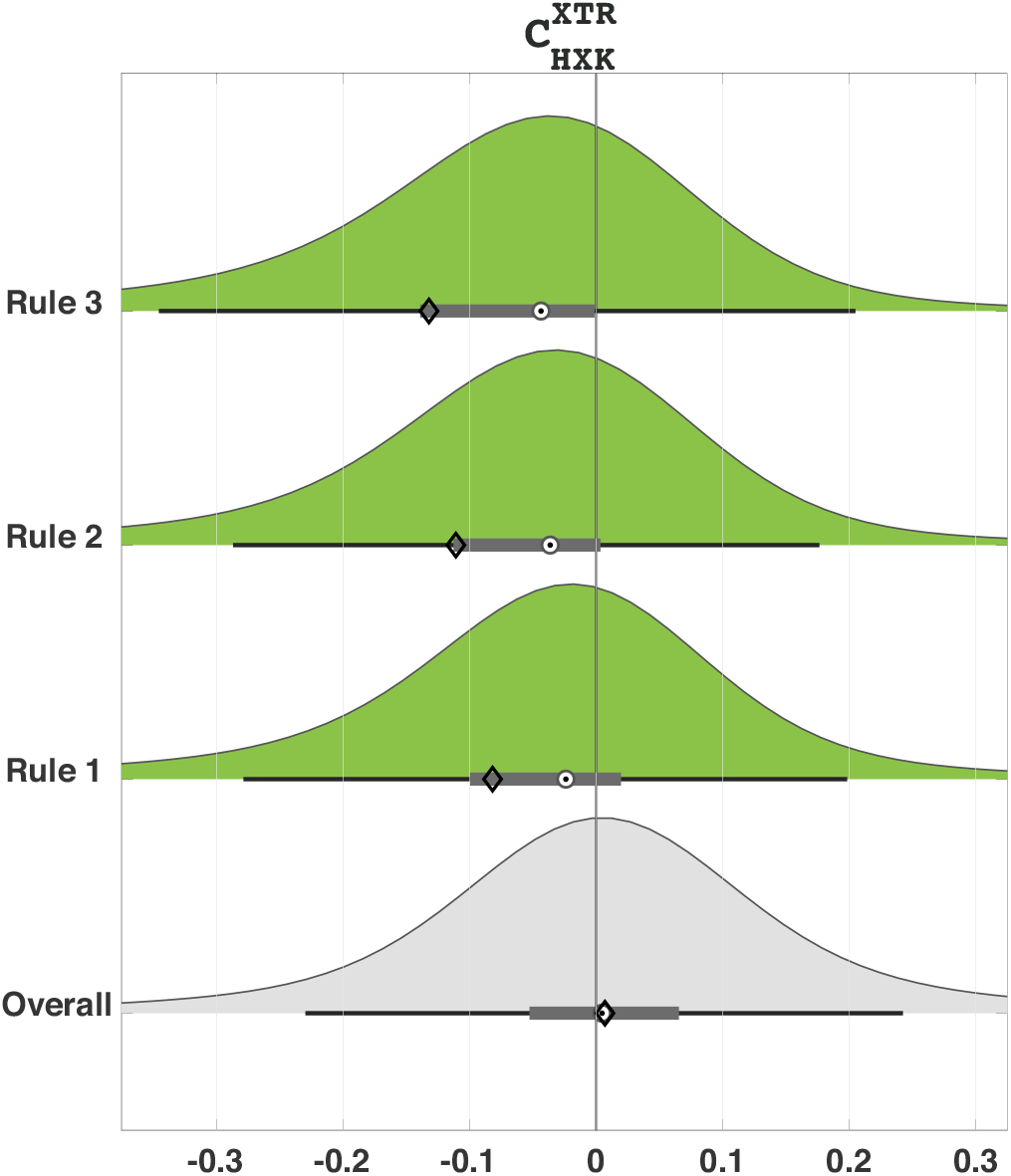
The distributions of control coefficients of XTR with respect to HXK for Rules 1, 2 and 3 were clearly shifted toward negative values compared to the one for the overall population of parameter sets. The horizontal box plots describe the interquartile range (gray box), median (target circle), mean (diamond) and the range of +/− 2.7 standard deviations around the mean (whiskers) of the distributions.

**Table 1:** Output of parameter classification algorithm for 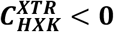. Top 3 rules obtained from the parameter classification with preselected parameters. The rules are ranked according the number of parameter sets that satisfy parameter ranges defined by the corresponding rule. For example, 9285 out of 200’000 (4.6%) generated parameter sets are within the subspace defined by the top-ranked rule, Rule 1. The values of σ_A_ relate to the *K_m_* values as *K_m_* = *S* (1 − σ_A_)/σ_A_, where *S* is the concentration of the corresponding metabolite. The notation 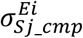 represents the degree of saturation of the enzyme *E_i_* by the metabolite *Sj*, and *cmp* denotes either cytosolic or mitochondrial compartment.

| Rule | Size | PI<br>($C_{HXK}^{XTR} < 0$) | Parameters | $K_m$ ranges (mM) | | $\sigma_A$ ranges | |
| --- | --- | --- | --- | --- | --- | --- | --- |
| 1 | 9285<br>(4.6%) | 0.65 | $\sigma_{pi\_m}^{ASN}$ | $3.5 \cdot 10^{-3}$ | $6.5 \cdot 10^0$ | 0.001 | 0.651 |
| | | | $\sigma_{t3p\_c}^{TPI}$ | $2.0 \cdot 10^{-5}$ | $2.2 \cdot 10^{-2}$ | 0.476 | 0.999 |
| | | | $\sigma_{dhap\_c}^{GPD1}$ | $7.1 \cdot 10^{-1}$ | $2.0 \cdot 10^3$ | 0.001 | 0.737 |
| | | | $\sigma_{g3p\_c}^{GPD2}$ | $5.1 \cdot 10^{-4}$ | $1.6 \cdot 10^1$ | 0.001 | 0.970 |
| | | | $\sigma_{adp\_c}^{ADK}$ | $3.4 \cdot 10^{-2}$ | $1.3 \cdot 10^2$ | 0.001 | 0.799 |
| | | | $\sigma_{glyc\_c}^{GLYct}$ | $9.9 \cdot 10^{-2}$ | $6.9 \cdot 10^2$ | 0.001 | 0.875 |
| | | | $\sigma_{nadh\_c}^{XRI}$ | $5.2 \cdot 10^{-2}$ | $7.6 \cdot 10^1$ | 0.001 | 0.593 |
| | | | $\sigma_{nad\_c}^{XRI}$ | $3.4 \cdot 10^{-4}$ | $2.6 \cdot 10^{-1}$ | 0.561 | 0.999 |
| 2 | 8049<br>(4.0%) | 0.73 | $\sigma_{adp\_m}^{ASN}$ | $2.6 \cdot 10^{-3}$ | $5.6 \cdot 10^0$ | 0.318 | 0.999 |
| | | | $\sigma_{pi\_m}^{ASN}$ | $6.5 \cdot 10^{-6}$ | $3.5 \cdot 10^{-3}$ | 0.651 | 0.999 |
| | | | $\sigma_{atp\_m}^{ASN}$ | $6.2 \cdot 10^{-3}$ | $1.6 \cdot 10^1$ | 0.001 | 0.719 |
| 3 | 6342<br>(3.2%) | 0.76 | $\sigma_{t3p\_c}^{TPI}$ | $2.0 \cdot 10^{-5}$ | $2.5 \cdot 10^{-2}$ | 0.443 | 0.999 |
| | | | $\sigma_{nadh\_c}^{XRI}$ | $1.1 \cdot 10^{-1}$ | $7.6 \cdot 10^{+1}$ | 0.001 | 0.418 |
| | | | $\sigma_{adp\_m}^{ASN}$ | $5.6 \cdot 10^0$ | $2.6 \cdot 10^{+3}$ | 0.001 | 0.318 |
| | | | $\sigma_{pi\_m}^{ASN}$ | $6.5 \cdot 10^{-6}$ | $3.5 \cdot 10^{-3}$ | 0.651 | 0.999 |
| | | | $\sigma_{atp\_m}^{ASN}$ | $6.2 \cdot 10^{-3}$ | $1.6 \cdot 10^{+1}$ | 0.001 | 0.719 |
| | | | $\sigma_{nadh\_c}^{XRI}$ | $2.8 \cdot 10^{-2}$ | $7.6 \cdot 10^{+1}$ | 0.001 | 0.733 |

#### Top 3 significant parameters

A closer inspection of the top ranked rules revealed that there were a few parameters such as 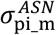 or 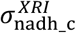 (for notation see the caption of Table 1) that appeared rather consistently throughout the rules (Table 1 and S2 File). The appearance of a reduced number of parameters throughout the inferred top ranked rules suggested that these parameters were essential for a negative control of HXK over XTR. We hence ranked the parameters based on the number of their occurrences in the rules and by how much their ranges were constrained (Methods).

We first considered the top rule (Rule 1 in Table 1 and S2 File), and we computed the ranking score for the associated parameters. We then ranked the parameters for the top 2 rules (Rules 1 and 2), for the top 3 rules (Rules 1, 2 and 3), and so forth, and observed how the ranking score of the parameters evolved as we considered a growing number of rules (Fig 3A). There was a clear separation in the ranking scores of a small number of parameters from the remaining parameters (Fig 3A). Indeed, the three highest ranked parameters, 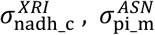, and 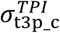, were consistent for a large number of considered rules. This result suggested that it would be sufficient to constrain a combination of the ranges of the three parameters to ensure the negative control of HXK over XTR.

**Fig 3.**
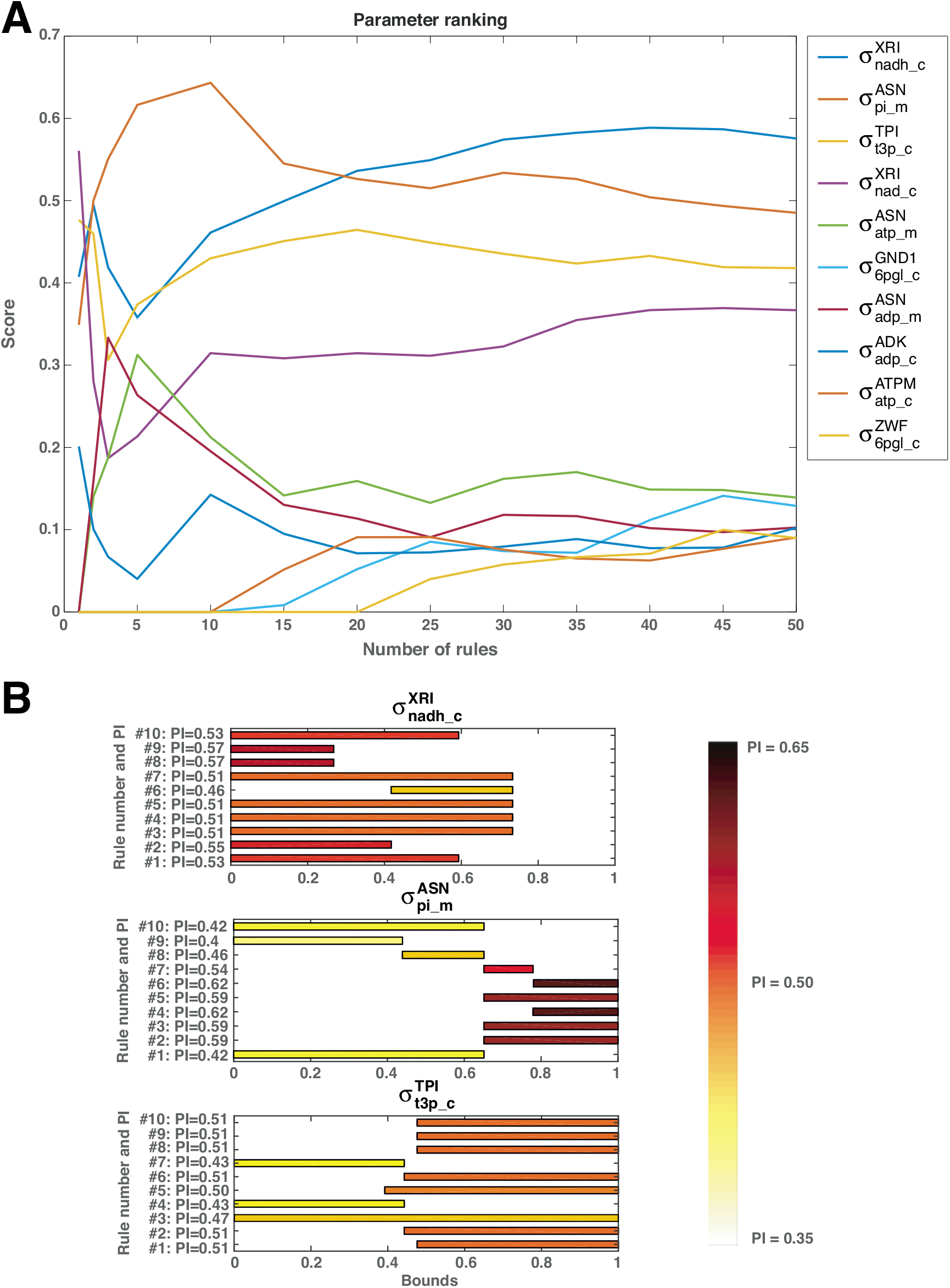
Top ranked parameters affecting control of hexokinase (HXK) over xylose uptake (XTR). **(A)** Evolution of the ranking score for the top 10 parameters as a function of the number of considered rules. **(B)** The effects of constraining the top 3 parameters individually according to the ranges of the top 10 rules on PI.

#### Qualitative dependency of negative control on top 3 significant parameters

We constructed a subspace of parameters by constraining the range of the top significant parameter, 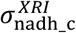, according to Rule 1 (Table 1), while the other parameters were unconstrained and could take any value between 0 and 1. Within this subspace, there were 53% of parameter sets giving rise to the negative control of HXK over XTR, i.e., PI=0.53 (Fig 3B, top). In such a way, we constrained the ranges of 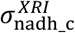 based on the remaining top 10 ranked rules, and we analyzed how these ranges affected PI (Fig 3B). There was a clear qualitative relationship between the ranges of 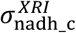 and PI. Indeed, the values of PI ranged from 0.55 (Rule 2) up to 0.57 (Rules 8 and 9) for low values of 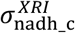, whereas they were as low as 0.46 for middle range values of this parameter (Fig 3B). We repeated this analysis for 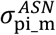 and 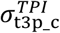, and for higher values of these two parameters, PI was as high as 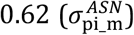 and 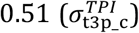, whereas for lower values of these two parameters PI was as low as 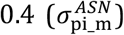 and 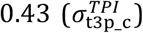. This observation motivated us to analyze how PI evolved with the progressive increase of the lower bound for each of the parameters while keeping their upper bounds at 1. Interestingly, the increase of the parameters lower bound lead to either a monotonic increase (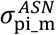 and 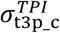) or decrease 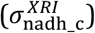 of PI (S4 Fig). Therefore, the effect of the parameters on the control of HXK over XTR was the most pronounced for the parameter ranges either in the low or the high values but not in the middle range.

This analysis suggested that a subspace defined by constraining 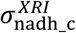 to low values and 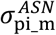 and 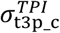 to high values was likely to have a high PI.

### Constraining top 3 significant parameters ensures the negative control of HXK over XTR

To combine the distributions of top 3 parameters that ensure a high PI in an unbiased way, we performed another parameter classification (Methods). The parameter classification algorithm inferred 66 rules on these three parameters, and the top rule enclosed 9389 samples with PI of 0.73 (S2 File). The PI value of 0.73 was close to the maximal PI value of 0.78, which was computed for the rules formed with all parameters. As expected, the ranges of the three parameters defined by the top rule (Fig 4C) were consistent with the analysis presented in the previous section.

**Fig 4.**
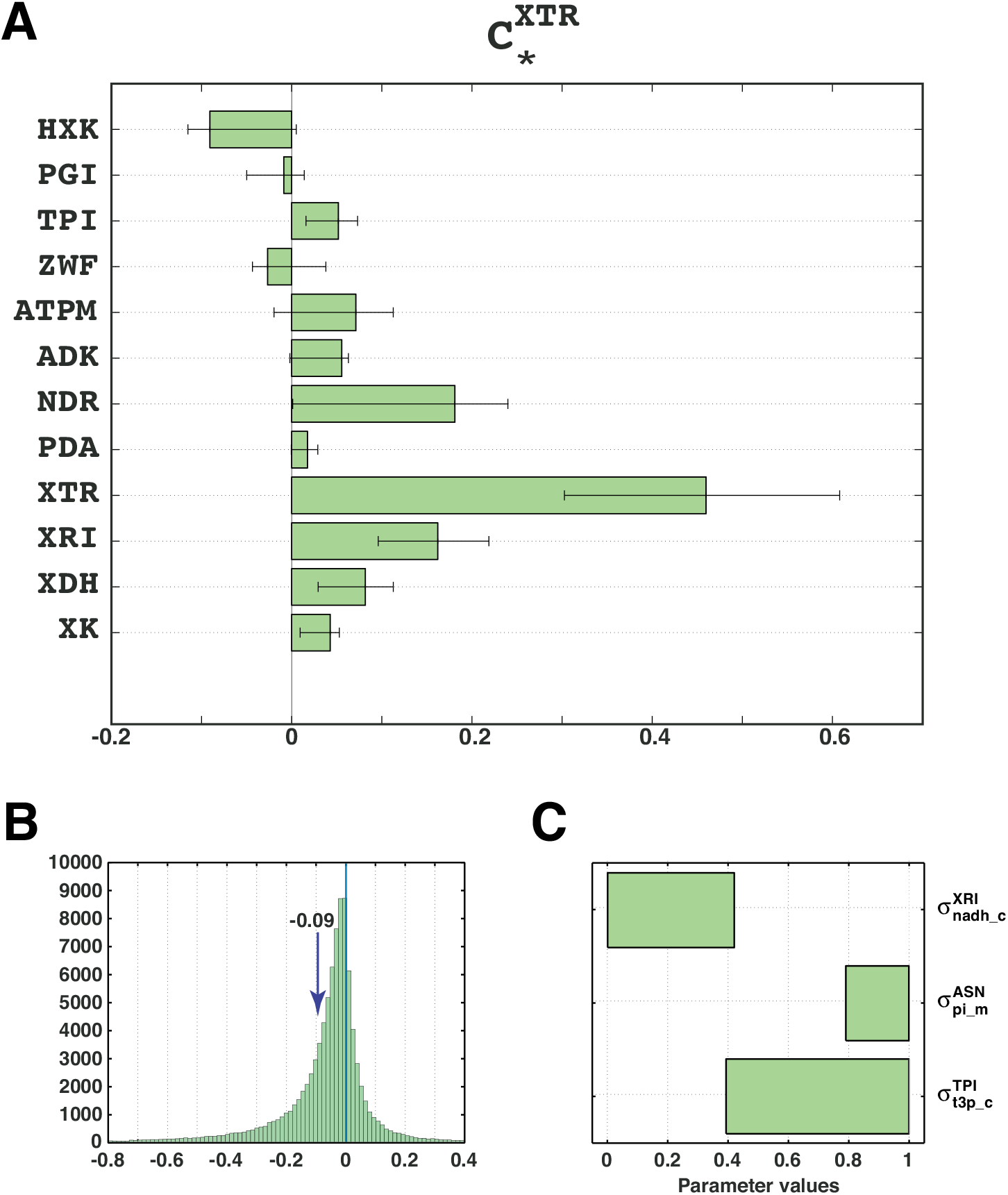
A well-determined control of HXK over XTR. **(A)** The mean control coefficient of HXK over XTR is negative after constraining the inferred ranges of only 3 parameters. **(B)** Distribution of the control coefficient 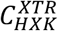. **(C)** The inferred ranges of 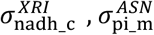, and 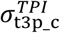 that determined the negative control of HXK over XTR.

We proceeded with the validation of the ranges of the top 3 parameters on a new population of models. We imposed the ranges of the top 3 parameters derived from the top rule of the parameter classification and generated a population of 100’000 models (Methods). We then computed the control coefficients of the top enzymes over XTR (Fig 4). The control coefficient 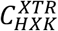 was distinctively negative with a mean value of −0.09 (Fig 4C), and its distribution was clearly shifted toward negative values compared to that of the original population of models (Fig 1B). More than 72% of models had negative values of 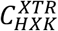 compared to 47% in the original population of models. The value of PI of 0.72 obtained from the validation set was strikingly close to the predicted value of 0.73 from the second tree training.

### Uncertainty in physiology: alternative concentration profiles

In Metabolic Control Analysis (MCA), it is considered that the control coefficients depend only on elasticities, however this holds only when the reactions are irreversible and there are no conserved moieties. It has been shown that metabolite concentrations affect displacements of reactions from thermodynamic equilibrium, which in turn influence the control over fluxes and concentrations in the network (3, 16, 32, 37). Therefore, when there is uncertainty in physiology, e.g., when several alternative concentration profiles correspond to experimental observations, the control coefficients derived from the kinetic models computed for these concentration profiles can be significantly different. iSCHRUNK can resolve this kind of problems by identifying the parameter values that give rise to well-determined control coefficients of the metabolic network for multiple alternative physiologies.

As an illustration, we analyzed three alternative physiologies characterized with three alternative concentration profiles (Reference, Extreme1 and Extreme2) and a common flux profile (Methods). We have undertaken to identify significant parameters that ensure a well-determined control over XTR for these physiologies. For this purpose, we constructed two populations of 200’000 kinetic models for the Extreme1 and Extreme2 physiology (Methods). Overall, together with 200’000 parameter sets computed previously for the reference physiology (Reference), we had 600’000 parameter sets for parameter classification. In the three populations of models, 47% (Reference), 46% (Extreme1) and 39% (Extreme2) of the models had a negative 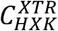.

#### Significant parameters for negative control of HXK over XTR for three alternative physiologies

For each of three populations of models, we preselected parameters based on the Fisher’s linear discriminant score, and we performed the parameter classification to find parameter ranges that guarantee a negative control of HXK over XTR (S2 File). We used the inferred rules from the three parameter classifications to rank the top parameters over the three alternative physiologies (Methods). Interestingly, the top seven parameters from the Reference case remained in the group of top seven parameters over the three cases (Fig 3 and S5 Fig). Moreover, the top two parameters (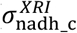 and 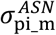) from the Reference case (Fig 3) were the top two also for the Extreme1 and Extreme2 case (S5 Fig). In contrast, 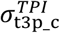 was less significant for the Extreme1 and Extreme2 case, and it was ranked below 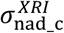 in the overall score (S5 Fig).

To refine the distributions of top 3 parameters for each of the three alternative physiologies, we performed three additional parameter classifications (Methods). However, the refined distributions of top parameters that ensure a negative control of HXT over XTR might or might not coincide for the three concentrations. Therefore, to reconcile the parameter distributions for the three cases, we used the parameter sets defined by the top 3 rules for each of these cases as input for an additional parameter classification (Methods). The top rule obtained from this parameter classification enclosed 11’801 out of 600’000 parameter sets, and 70.9% of these models had a negative control of HXK over XTR (Table 2).

**Table 2:**
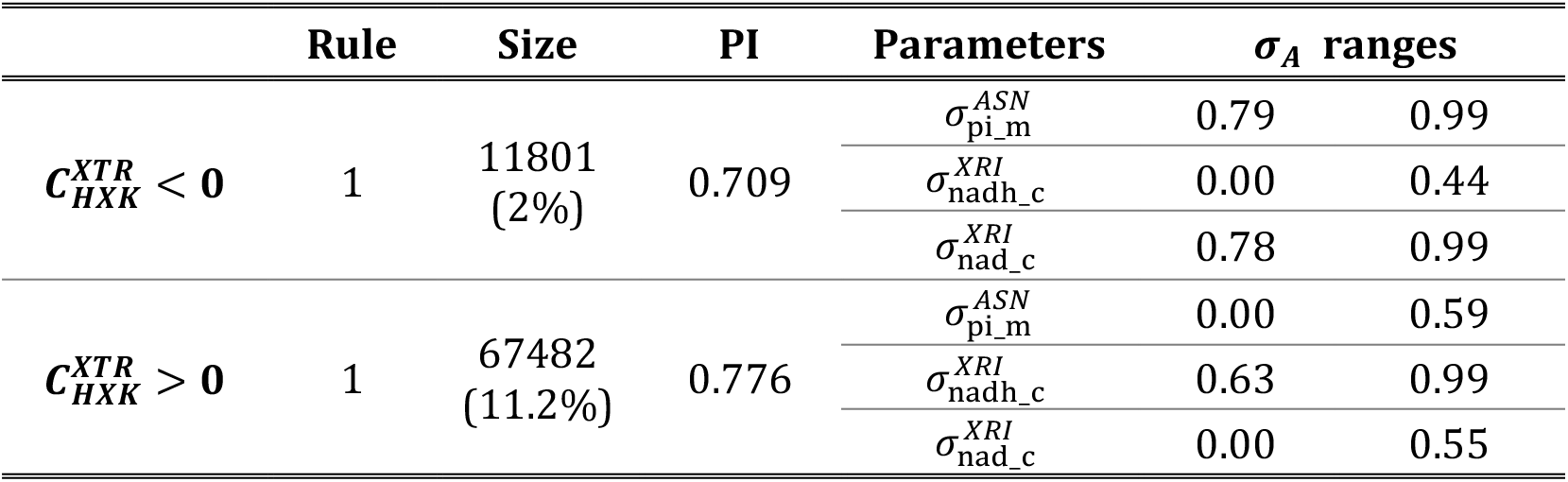
Robust ranges of the top 3 parameters over 3 concentrations.

We imposed the robust ranges for the top 3 parameters from Table 2 and generated a population of 100’000 models for each of the three alternative physiologies, a total of 300’000 models. We then computed the control coefficients of HXK over XTR, and a significant improvement of PI was obtained for all three alternative physiologies. Indeed, the Reference, the Extreme1 and the Extreme2 cases, had 72%, 72% and 67% of models with a negative 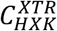, respectively (compared to the 47%, 46% and 39% of models with unconstrained parameters). The average PI (0.703) over three populations of models was remarkably close to the predicted PI (0.709) (Table 2).

#### Significant parameters for positive control of HXK over XTR for three alternative physiologies

We repeated the procedure from the previous section using the same set of top 3 parameters, but we imposed a positive control of HXK over XTR as the objective for the parameter classification. The top inferred rule enclosed a significantly higher number of models (67482) compared to the case with a negative control of HXK over XTR (11801) (Table 2). Assuming that the parameter space was sampled uniformly, this also suggested that the parameter subspace that ensures a positive control of HXK over XTR was larger than the one that ensures a negative control. Interestingly, there were no overlaps between parameter ranges for a positive and a negative control (Table 2). The predicted PI for a positive control (0.776) was also higher than the one predicted (0.709) for the negative control.

We generated a population of 100’000 models for each of the three alternative physiologies by imposing the robust distributions of the top 3 parameters ensuring a positive control (Table 2). We obtained PI of 0.747 (Reference), 0.802 (Extreme1), and 0.767 (Extreme2) (Table 3), which was a notable improvement compared to the population with unconstrained parameters with PI of 0.53 (Reference), 0.54 (Extreme1), and 0.61 (Extreme2). Similar to a negative control study, the average PI over three populations of models (0.772) was remarkably close to the predicted one (0.776) (Table 3).

**Table 3:** Validation of the robust distributions of the top 3 parameters for both a negative and a positive control of HXK over XTR.

|  | Reference<br>physiology | Extreme 1<br>physiology | Extreme 2<br>physiology | Average |
| --- | --- | --- | --- | --- |
| **PI**<br>**($C_{HXK}^{XTR} < 0$)** | 0.719 | 0.720 | 0.669 | 0.703 |
| **PI**<br>**($C_{HXK}^{XTR} > 0$)** | 0.747 | 0.802 | 0.767 | 0.772 |

These results showed that few parameters determine whether the control of HXT over XTR is negative or positive, meaning that the operational states of few enzymes are vital to responses of the metabolic network upon perturbations. For example, a follow-up experiment testing ambiguous control of HXK over XTR was performed in (30), and it was shown that *HXK2* deletion improves xylose uptake rate. Based on this experimental observation, one can hypothesize the operating ranges of top enzymes such as ATP synthase (ASN), triose phosphate isomerase (TPI) or xylose reductase (XRI).

The obtained parameter values of kinetic models pertaining to single physiology, e.g., Reference or Extreme1 had both higher PIs and larger parameter subspaces compared to the ones inferred from models constructed for the three alternative physiologies. The reason behind this is that different metabolite concentrations result in different operating ranges of enzymes, thus affecting the control over the analyzed quantities. One might expect that the distributions of parameters ensuring a high PI will shift, and possibly shrink or stretch, as concentration values change. Thus, to obtain values of the studied parameters that ensure a high PI over multiple concentrations, we combined ranges of parameters computed for individual concentrations and curated them through a parameter classification (Methods).

We computed here robust distributions of parameters for only three, albeit very different, metabolite concentration vectors, nevertheless, the proposed method can readily be used to find parameter distributions for a large number of concentration vectors. Thus, iSCHRUNK can be used to create a mapping between the metabolite concentrations and kinetic parameters spaces, and identify the regions in the parameters, metabolite concentrations, and thermodynamic displacements from equilibrium spaces that give rise to a systemic property, e.g., robust steady-state responses to genetic and environmental perturbations.

### Improved parameter classification through reassignment

We found no rule with PI equal to 1 in performed parameter classification studies. This suggested that the parameter subspaces leading to a negative and a positive control of HXK over XTR were not distinctly separated. To improve the parameter classification for the problems where the separation between the classes is fuzzy, we propose to employ the *k*-nearest neighbors (*k*-NN) algorithm (Methods). The *k*-NN algorithm allows us to identify the parameter sets from one class that are surrounded by the parameter sets of the other class and reassign them to the latter. In the context of finding parameter values that give rise to a certain property, this means that the parameter classification algorithm will find only those parameter sets that are surrounded by a majority of the parameter sets of the same class. This way, the separation between the classes will be increased at the expense of neglecting parameter sets from the regions with a heavy overlap of the classes.

We reconsidered the classification for parameters determining a negative control of HXK over XTR in the Reference case, and we applied the *k*-NN algorithm with *k*=10 over the set of initial 200’000 parameters in order to find the surrounding for each of parameter sets, and to perform the reassignment (Methods). If in the group of 10 closest neighbors of a parameter set the percentage of parameter sets from the same class was less than a reassignment threshold, *r*, then that parameter set was reassigned. We performed two parameter classification studies for two different reassignment thresholds, *r*, of 30% and 50% (Methods).

We found that as the reassignment threshold was increasing the tree training algorithm was inferring a smaller number of rules (73 for *r* = 30% versus 31 for *r* = 50%). Furthermore, the inferred rules were enclosing a smaller number of parameter sets for higher values of *r*, i.e., for *r* = 30% and 50%, the top rules enclosed respectively 13427 and 1339 parameter sets (Fig 5 and S2 File). In contrast, the obtained PIs, were higher for *r* = 50% than for *r* = 30% (Fig 5). For example, PI of the top rule for *r* = 50% was 0.83, whereas the one for *r* = 30% was 0.73 (Fig 5 and S2 File). A comparison between the original method with preselection, which is identical to the reassignment method with *r* = 0% (corresponding to no reassignment), and the reassignment methods for *r* = 30% and 50% showed a general tendency of the latter for obtaining rules with improved PI and that enclose a smaller number of parameter sets (Fig 5).

**Fig 5.**
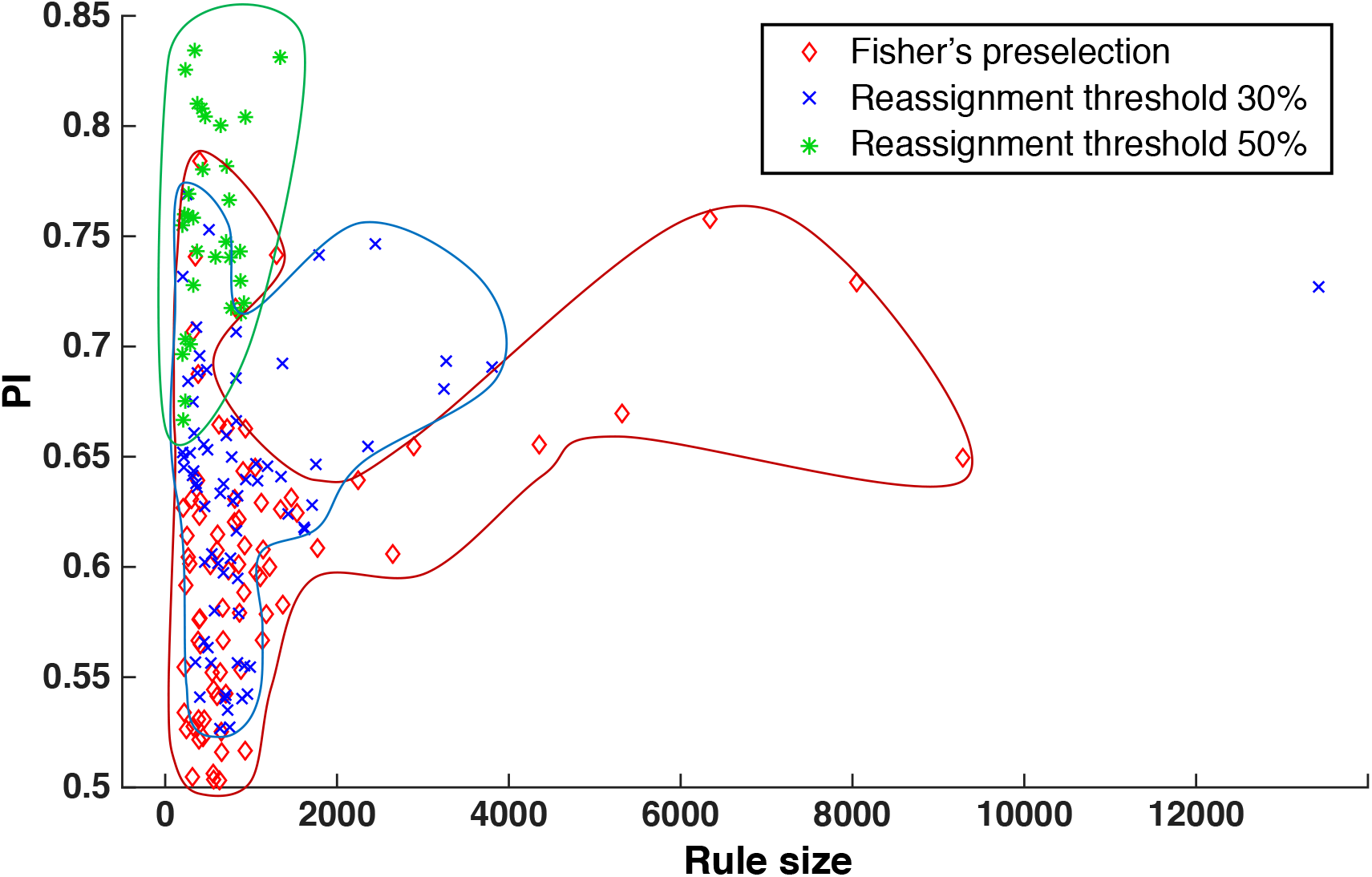
The rules obtained with the original method versus the ones obtained with the improved parameter classification algorithm. For each rule, we plot the performance index (PI) as a function of the number of enclosed parameter sets (rule size). The rules are obtained with: the original method with preselection (red diamonds), the reassignment method with 30% threshold (blue crosses) and the reassignment method with 50% threshold (green asterisks).

We also tested the reassignment procedure for parameters determining a positive control of HXK over XTR in the case of the reference metabolite concentration with *k*=10 and *r* = 60%. The classification algorithm inferred 19 rules with PIs ranging from 0.75 to 0.90.

The rules were defined by only 28 parameters (S2 File). The top rule enclosed 1711 parameter sets with PI of 0.90, and it was defined by 6 parameters.

To validate the proposed improvement to the parameter classification, we imposed the distributions of the parameters defined by the top rules for the negative control case with r=50%, and for the positive control case with r=60% (S2 File). We generated for each study a population of 100’000 models, and we computed the control coefficients in the network.

In the case of negative control, the distribution of the control coefficient 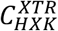 was biased toward negative values with mean −0.13 (Fig 6A and 6B). More than 79% of the computed control coefficients 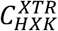 were negative. (Table 4). Similarly, in the case of positive control, the distribution of the control coefficient 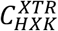 was shifted toward positive values with the mean of 0.21 and a remarkable PI of 0.89 (Fig 6D and 6F, and Table 4). For the negative and positive cases, the top rules were defined by 6 parameters each, where three parameters, 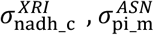, and 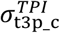, were common for both cases (Fig 6C and 6E). These three parameters were also ranked as the top 3 parameters in the parameter classification with the original algorithm (Fig 4). Moreover, the range of 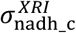 was constrained toward low values for the negative control and toward high values for the positive control. In contrast, the parameters 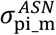 and 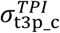 were constrained toward high values for the negative control, and toward low values for the positive control. These patterns suggest that these three parameters are crucial for determining the sign of the control of HXK over XTR, whereas the remaining parameters, 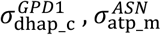, and 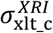 for the negative control case, and 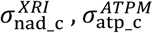, and 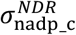 for the positive control case, are likely having a minor effect on the PI.

**Fig 6.**
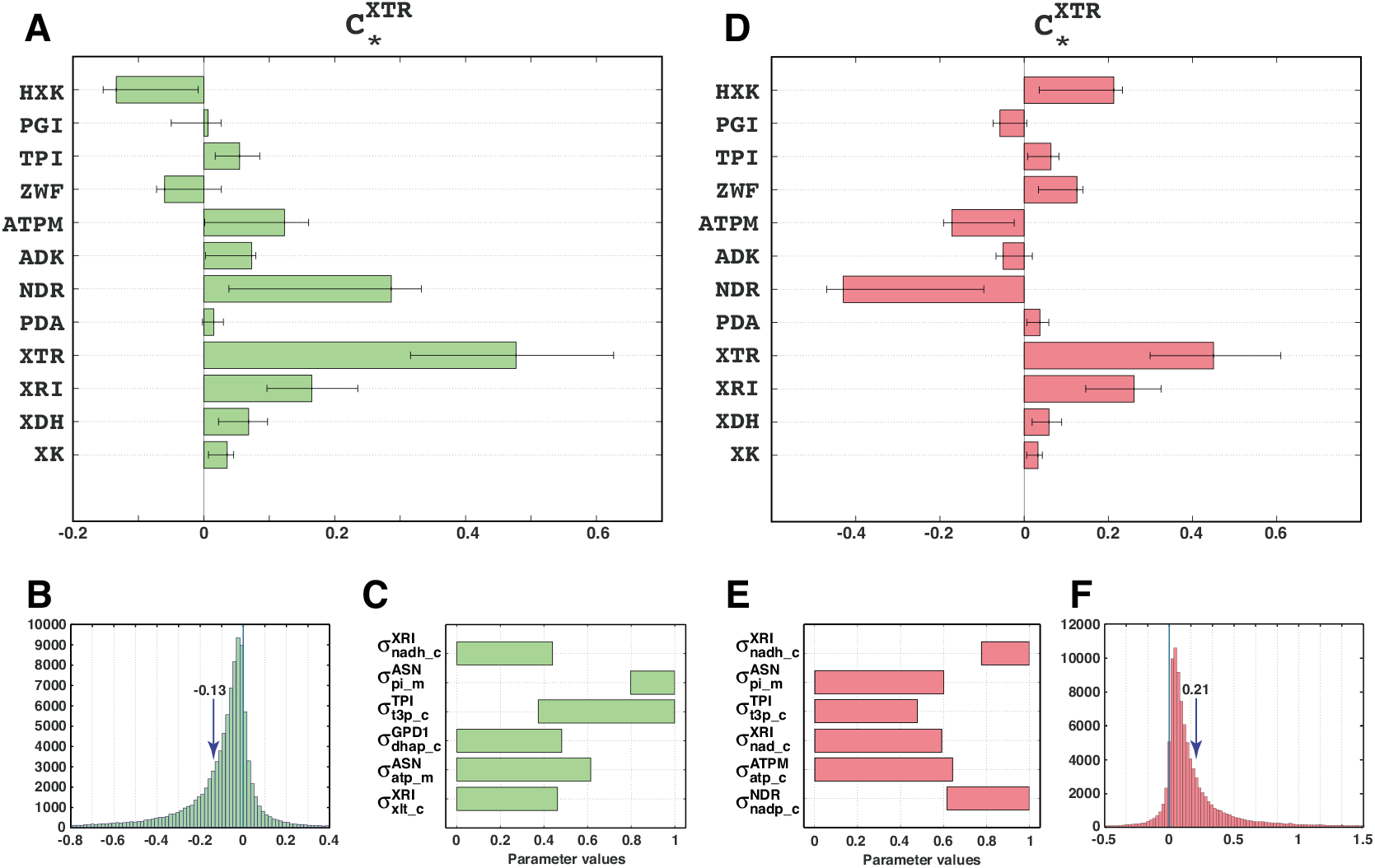
Validation of the parameter classification with reassignment. Control of HXK over XTR is negative **(A)** and positive **(D)** after constraining the inferred ranges of 6 parameters. Distribution of the control coefficient 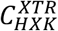 for the case of negative **(B)** and positive **(E)** control. The inferred ranges of parameters that determined negative **(C)** and positive **(F)** control of HXK over XTR.

**Table 4:**
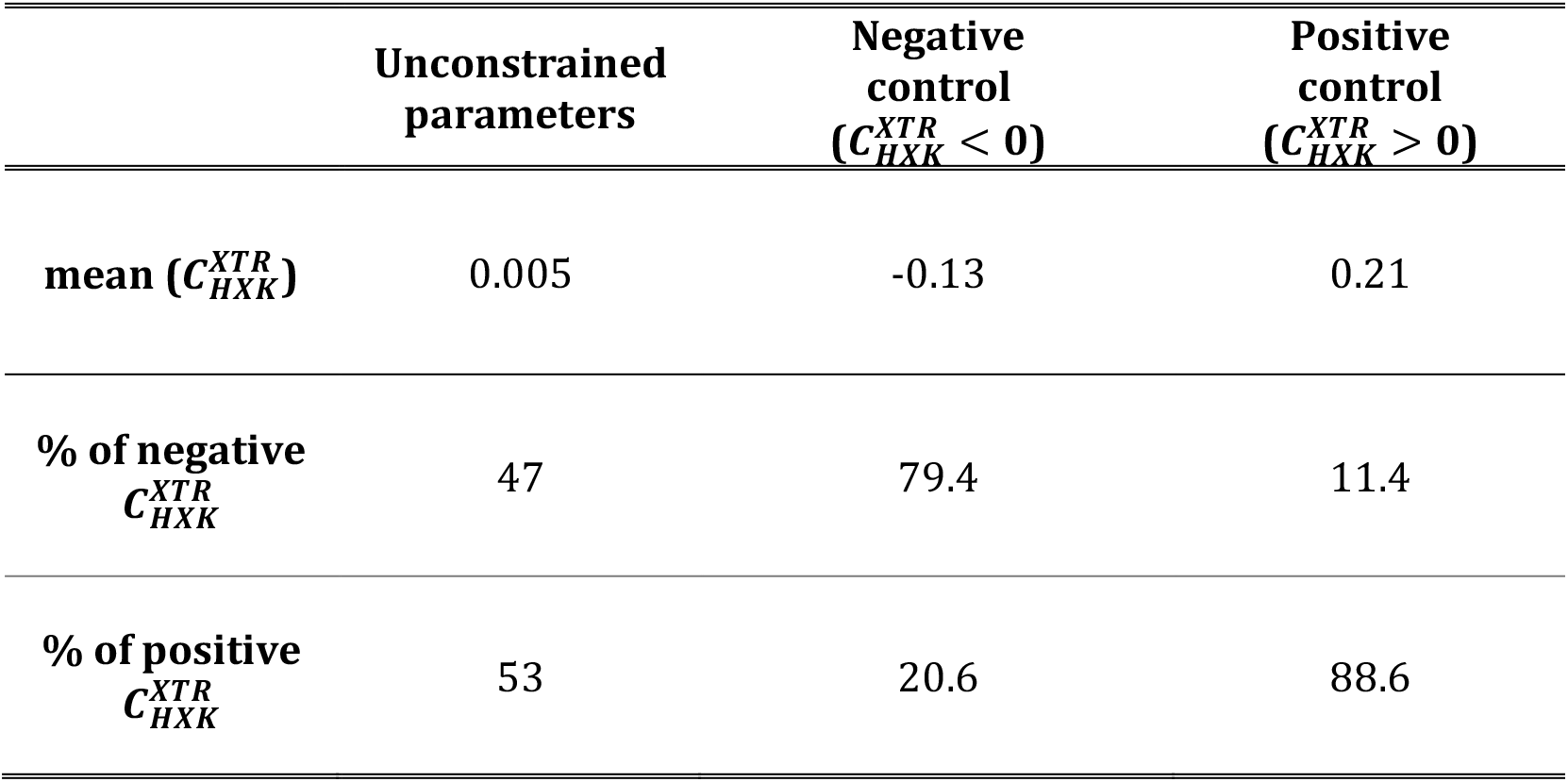
Validation of the ranges of the top 6 parameters obtained with reassignment for both a negative and a positive control of HXK over XTR.

This result clearly demonstrated that the reassignment procedure allows for more precise identification of the subspaces leading to a desired control of HXK over XTR. We observed improvement of both PI and the mean 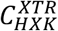 value compared to the results obtained with the unaltered parameter classification algorithm.

## Discussion

Machine learning methods (38–42) have found applications in a large number of biological and biomedical areas such as cancer research (43–45), population genetics (46, 47), protein structure and function prediction and phylogenomic mapping (48–52), protein-protein interactions (53–55), medical imaging (56–60), gene expression and microarray data analysis (61–64), regulatory interactions (65, 66), metabolic pathway dynamics (67), biomarker discovery and analysis of metabolomics and proteomics data (68–71). However, the potential of these methods for detecting patterns in parameters of kinetic models of metabolism and uncovering hidden relationships between kinetic parameters, omics data, and observed phenotypes remained largely unexploited.

Machine learning methods require large sets of training data for their successful application and methods for generating kinetic metabolic models that use Monte Carlo sampling offer an unprecedented opportunity for employing machine learning to advance our understanding of metabolic processes in cellular organisms. Kinetic models are usually built around a metabolic steady-state, which is characterized by the metabolite concentrations and metabolic fluxes, and the generated populations of kinetic parameters together with the observed steady-state data contain implicit information about the studied physiology. This information, if extracted systematically, can be used as guidance for the design of metabolic engineering and synthetic biology strategies that ensure the desired metabolic responses of studied organisms.

In this work, we have extended iSCHRUNK functionalities to data-mine this information and systematically reduce uncertainties in the values of kinetic parameters that give rise to the desired metabolic behavior. As a demonstration, we reduced the uncertainties in the kinetic parameters that ensure that values of flux control coefficients remain within a pre-specified range.

iSCHRUNK lends itself to a broad scope of applications ranging from sustainable production of biochemicals to medicine and regarding both the analysis and design of metabolism. It allows us to analyze the relationships between the inferred parameter ranges and the measurements acquired on the actual biological system, and, consequently, to create hypotheses regarding the operating states of enzymes and provide information about saturations of all enzymes in the network. This information is crucial for biotechnology studies where living cells need to be engineered for improved performance, or for drug discovery studies where, e.g., we want to overproduce a compound that is toxic to a pathogen.

The method can be applied not only to identify distributions of kinetic parameters but also to determine distributions of the metabolic fluxes and metabolite concentrations satisfying given requirements. It can also be used for guaranteeing both qualitative and quantitative features of metabolism, and several requirements can be combined simultaneously. For example, iSCHRUNK can be used to identify and quantify the parameters that maintain a redox potential while enforcing the desired level of yield and specific productivity of a compound of interest. Provided that the desired properties are biologically feasible, the method can be used to guarantee an arbitrary number of requirements.

Finally, iSCHRUNK can be used to alleviate issues with high computational requirements of Monte Carlo sampling of kinetic parameters in large- and genome-scale metabolic networks. As the size of the models and complexity of studies increases, sampling a kinetic space becomes increasingly difficult and even intractable. However, iSCHRUNK allows us to identify relevant kinetic parameters that correspond to the observed physiology. The key finding of the current and previous studies (17) is that only a small set of parameters corresponding to a few enzymes is sufficient to characterize the observed physiology. Therefore, once we identify the most relevant parameters, it suffices to densely sample the identified parameters while fixing the remaining parameters at arbitrary feasible values. This way, iSCHRUNK dramatically reduces the sampling space, thus enabling computational analyses of large-scale and genome-scale dynamic metabolic systems.

## Methods

### Identifying significant parameters that determine studied properties

The computational method for characterization and reduction of uncertainty, iSCHRUNK, was proposed in (17). iSCHRUNK involves a set of successive computational procedures that can help us to ascertain and quantify the kinetic parameters that correspond to a given physiology. iSCHRUNK can be used with any method that generates populations of kinetic models describing given physiology such as ensemble modeling (24) or ORACLE (3, 4, 8, 10, 11, 31, 32). Here, we extended the original iSCHRUNK workflow (17) by an iterative loop that uses parameter classification to perform stratified sampling of the kinetic parameters, i.e., it allows identifying refined sets of parameters that lead to the desired metabolic behavior (Fig. 7). We used the extended iSCHRUNK to identify the distribution of kinetic parameters that determine the sign in ambiguous distributions of control coefficients as follows:

I. We defined the stoichiometric model of glucose-xylose co-utilizing *S. cerevisiae* (S6 Fig). The model consisted of 102 atomically balanced reactions and 96 intracellular metabolites, and included glycolysis, pentose phosphate pathway, tricarboxylic cycle (TCA), electron transport chain (ETC) and XR/XDH xylose assimilation pathway (30, 72). Based on the physiological information on the cellular compartmentalization the intracellular metabolites were categorized as cytosolic or mitochondrial, and the extracellular metabolites were modeled as well. We integrated the thermodynamic constraints based on the information about the Gibbs free energies of reactions (73–76) together with the fermentation data from Miskovic et al. (30), and we then performed the Thermodynamics-based Flux balance Analysis (TFA) (4, 13–15, 77, 78) to compute the thermodynamically consistent steady-state flux (S7 File).
II. We sampled the space of metabolite concentrations that is consistent with: (i) the directionalities of the steady-state flux obtained in step I; and (ii) the available observations of metabolite concentration ranges (4, 8). The displacements of the reactions from thermodynamic equilibrium that correspond to the sampled metabolite concentration sets were simultaneously computed (18, 27). We then computed *the reference vector of metabolite concentrations*, Reference, as the sample that was closest to the mean metabolite concentration vector (S7 File). The Principal Component Analysis (4, 79) of the samples was next performed, and we computed *two extreme metabolite concentration vectors*, Extreme1 and Extreme2, as the two samples that were at the extreme ends of the sampled space along the direction of the first principal component (S7 File). In the following steps, we have computed a population of kinetic models for each of three alternative physiologies. The three physiologies were characterized by the common flux profile computed in Step I and three alternative concentration profiles (Reference, Extreme1 and Extreme2) computed in this step.
III. We assigned a kinetic mechanism to each enzyme-catalyzed reaction using the information from literature (18, 80–82). For reactions without available information about their kinetic mechanisms, we used the generalized reversible Hill law (83). The used kinetic mechanisms included reversible Michaelis-Menten kinetics, Uni-Bi, Bi-Uni, ordered Bi-Bi, Bi-Ter, Ter-Bi (81). We also modeled an allosteric regulation for the phosphofructokinase (PFK), where the assigned kinetic mechanism was Hill kinetics with the Hill coefficient *h* = 4 (S8-S10 File). At this point of the procedure, we may integrate available Michaelis constants, *K_m_*, from the literature and databases (84, 85). In this study, we did not use *K_m_* values from the literature, instead, we sampled the space of *K_m_* values indirectly through the sampling of the degree of saturation of the enzyme active site, σ_A_ (10). For a vector of metabolite concentrations computed in Step II, we calculated the *K_m_* value corresponding to a value σ_A_ as *K_m_* = S_j_(1 − σ_A_)/σ_A_, where S_j_ is the j^th^ element of the metabolite concentration vector that corresponded to σ_A_ (10). Without prior information, we sampled σ_A_ values between 0 (non-saturation) and 1 (full saturation). Otherwise, we performed the stratified sampling where we imposed the σ_A_ distributions obtained from the classification algorithm in Step VII (Fig 7A and 7B). An alternative to sampling σ_A_ values would be to sample the enzyme states (27, 28).
IV. We verified the local stability of the steady-state (10), and we rejected the kinetic parameters corresponding to unstable steady states and the ones that are not consistent with the experimentally observed data and literature.
V. In this step, we analyze whether or not the studied property is satisfied. If yes then we proceed to step VII, otherwise we perform the parameter classification in Step VI to find parameter values that give rise to the studied property. Here, we computed populations of control coefficients to quantify the responses of the metabolic fluxes and intracellular metabolite concentrations to changes in activities of the network enzymes. In general, we can study any property related to metabolic network such as: significant fluxes in the network such as the product flux and the uptake fluxes, yields, key concentrations such as ATP or NADH, other relevant quantities such as redox potential (NADH/NAD^+^). We then verified if the control of HXK over XTR was well determined. We defined the control of an enzyme over the analyzed quantity as being well determined if 50% of control coefficients around the mean control coefficient had the same sign. For example, in the population of the control coefficients of XTR with respect to xylulokinase (XK) all the samples between the 1^st^ and the 3^rd^ quartile were negative (Fig 1A), and hence we considered that XK had a well-determined negative control over XTR. In contrast HXK, ATPM and NDR had an ambiguous control over XTR (Fig 1A). If HXK had well-determined control over XTR, we proceeded to Step VII. Otherwise, we went to Step VI.
VI. We fed back to the classification algorithm the population of the analyzed control coefficient from Step V together with the corresponding values of the parameters (degree of saturation of the enzyme active site σ_A_) from Step III. The classification problem was defined to find the ranges of the σ_A_ values (and consequently the ranges of the corresponding *K_m_* values) that determine the sign, positive or negative, of the analyzed control coefficient. We solved this parameter classification problem using the CART algorithm (33, 34) from the MATLAB software package. We then used the output of the parameter classification, the distributions of σ_A_, for the sampling in Step IIIb (Fig 7A). More details about the parameter classification are presented in the next section.
VII. In this step, we can postulate hypotheses and design systems biology strategies.

**Fig 7.**
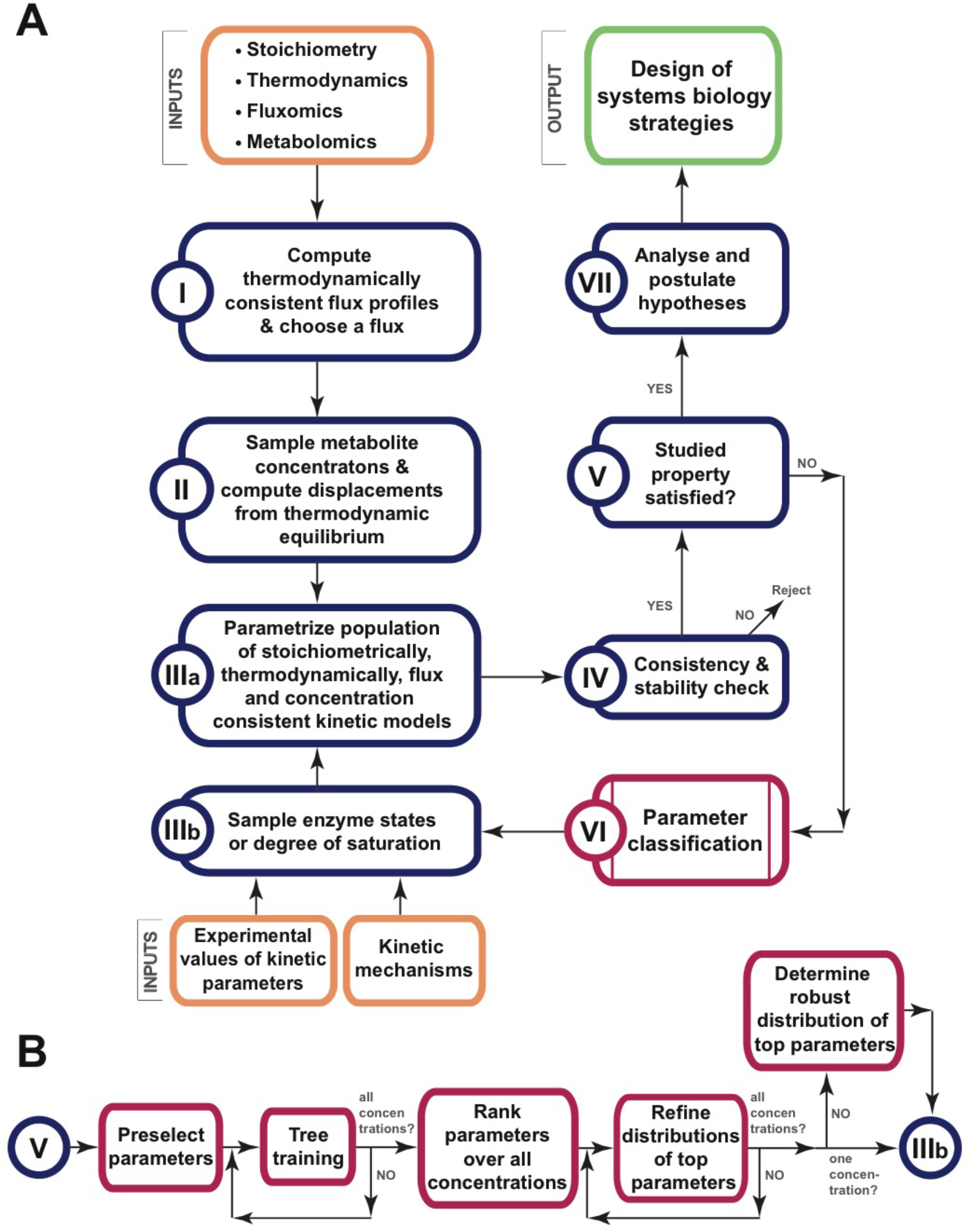
Workflow of the computational procedure for uncertainty analysis. **(A)** The workflow allows us to identify ranges of kinetic parameters ensuring that a studied property is satisfied, e.g., the sign in ambiguous distributions of control coefficients. **(B)** Detailed steps of parameter classification.

In Step V we entered an iterative loop for identifying the ranges of σ_A_ (or equivalently *K_m_*) for which the analyzed control coefficients were well determined (Fig 7). The iteration started by passing the invalidated σ_A_ values from this step to the classification algorithm in Step VI. We then used the refined σ_A_ distributions from Step VI in the sampling procedure in Step III. Next, the refined samples of σ_A_ were next tested for consistency in Step IV, and finally, we constructed a new population of control coefficients in Step V and verified it. At each iteration, the σ_A_ values (*K_m_* values) that reduced the ambiguity in the population of the analyzed control coefficients were refined and used for stratified sampling in Step III.

### Parameter classification

We carried out the parameter classification in several steps (Fig 7B). We first removed from the consideration the parameters that were not affecting the control over the analyzed flux. We then used the CART algorithm with the preselected parameters for three populations of kinetic models where each population was computed with a different metabolite concentration vector (see Step II of the framework discussed above). In the third step, we ranked the parameters over three concentrations, and we chose the top parameters to continue. We next refined the distributions of the top parameters for each concentration individually, and we then used this information to determine the consistent distributions of top parameters over all concentrations. We detail the parameter classification steps below.

#### Preselect parameters

Our preliminary results indicated that only a subset of kinetic parameters affected the sign of the analyzed control coefficient. The reduction in the parameter space was in agreement with our previous study (17), and inspired us to assess which parameters had a negligible effect on the computed control coefficients, to discard them, and then to proceed with the parameter classification. The benefits of preselecting the parameters are twofold. First, applying a computationally inexpensive method for preselecting the parameters and then using the CART algorithm on the reduced space can significantly reduce computational requirements of iSCHRUNK. Second, the parameters with a negligible effect on the control coefficients can introduce a bias in the estimates of key parameters. We can eliminate this bias by discarding the irrelevant parameters. We used the Fisher’s Linear Discriminant score (35, 36) to preselect the parameters:

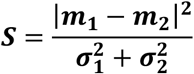

where *m*_1_ and *m*_2_ denote the mean values of the parameter populations that result in negative (or positive) and non-negative (or non-positive) control coefficients, while 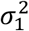 and 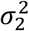 are the corresponding variances. The higher *S* was, the larger was the influence of the analyzed parameter in discriminating between a positive and a negative control. We ranked the parameters according to this score, and we kept the parameters whose scores were at least 1% of the highest obtained score.

#### Train classification tree and rank inferred classification rules

For each of the three metabolite concentration vectors, we trained a classification tree. The classification algorithm inferred classification rules based on the values of the preselected σ_A_ parameters and the outcomes, e.g., negative and non-negative control coefficients, obtained with these σ_A_ parameters. Each rule corresponds to a set of inequalities defined on different parameters. Thus, a rule is a hypercube in the parameter space. The number of inferred rules depends on the properties of the parameter space to be classified and also on the number of parameter sets that are used to train the algorithm. In order to prevent the overfitting, we fixed to 200 the minimal number of parameter sets that the algorithm can use to construct a rule (17, 34, 86). The rules defined by a large number of parameter sets are “more certain”. Besides, assuming that we sampled the parameter space uniformly, the “more certain” rules will likely enclose a larger volume of the parameter space with the well-determined control. Therefore, for each metabolite concentration vector, we ranked the inferred classification rules according to the number of parameter sets they contained.

#### Rank parameters across classification rules and over all concentrations

To rank the parameters of the models obtained for a concentration vector such as reference concentration vector or one of the extreme concentration vectors, we defined the following score for parameter *j*:

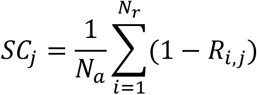

where *N_a_* denotes the number of rules over which we performed the ranking, *N_r_* denotes the number of rules, a subset of *N_a_* rules, that constrained parameter *j*, and *R_i,j_* is the range of parameter *j* defined by the rule *i*. This score incorporates two factors: (i) a number of occurrences of parameters across classification rules – parameters that appear in more rules are more relevant; (ii) how much parameters are constrained – less important parameters are less constrained.

To rank the parameters of the models obtained over all concentrations, we computed the aggregate score:

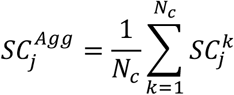

where *N_c_* denotes the number of metabolite concentration vectors and 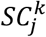 is the score computed for the concentration *k*. Observe that values of *N_a_, N_r_* and *R_i, j_* may differ for different concentrations.

We can choose to perform the ranking across: (i) all rules returned by the classification algorithm; though the classification algorithm can return different number of rules for three metabolite concentrations, the normalization constants *N_a_* ensure an unbiased scoring over different concentrations; (ii) the chosen top rules, e.g., over Top 10 rules (*N_a_* = 10 for all three concentrations). We used this score to rank the parameters; the higher the score was for a parameter, the higher was its ranking.

#### Refine distributions of top parameters

For each of metabolite concentration vectors (Reference, Extreme1 and Extreme2) and a set of top parameters ranked over all concentrations, e.g., Top 3 or Top 5 σ_A_ values, we performed the second parameter classification, i.e., tree training, to find refined parameter distributions that determine the sign of the analyzed control coefficient. The inputs to the parameter classification algorithms were the population of the analyzed control coefficient together with the corresponding top σ_A_ values. Thus, for the three concentration vectors, we constructed parameter subspaces that were constrained only by the obtained ranges of the top parameters.

#### Determine robust distributions of top parameters over alternative physiologies

The refined distributions of top parameters that correspond to a well-determined control might well mismatch among the three cases (Reference, Extreme1 and Extreme2). Therefore, some parameter values can correspond to a well-determined control for one physiology and to an ambiguous control for the other physiologies. To obtain the consistent distributions of the top parameters over all concentrations in an unbiased way we performed the third tree training as follows.

For each of the three alternative physiologies, we took as the input to parameter classification the parameter sets whose ranges of top parameters were defined according to top 3 rules. We used in parameter classification the parameter sets from the subspace of top rules as these parameter sets are likely to have well-determined control at least for one of the alternative physiologies. We then verified for each top parameter set if it corresponds to a well-determined control for the three alternative physiologies. If a parameter set corresponded to a well-determined control for the three alternative physiologies (S11 Fig, red stars), we considered it consistent; otherwise, it was considered inconsistent (S11 Fig, blue and yellow stars). We fed this information as the second input to the classification algorithm and performed the training.

The obtained consistent distributions of top parameters over the three cases were used in Step III to perform a stratified sampling.

### Reassignment procedure for improved tree training

In the cases when the space of parameter sets leading to a negative and the one leading to a positive control over analyzed quantities are overlapping, the separation between parameter classes is fuzzy. To enhance the separation between the classes, we propose here utilization of the *k*-nearest neighbors (*k*-NN) algorithm in the parameter classification as follows (87).

For each of the parameter vectors, we first assessed whether or not they were determining, e.g., a negative control, and we assigned them to two distinct sets. The first set, *SN*, contained parameter vectors that gave rise to a negative control, whereas the second set, *SP*, contained the ones that gave rise to a non-negative control. We then ran the *k*-nearest neighbors (*k*-NN) algorithm, and for each parameter vector from the set *SN*, we computed how many out of its *k*-nearest neighbors belonged to the same set (SN). For each of these parameter vectors, if the percentage of *k*-nearest neighbors that belonged to the set *SN* was higher than a pre-specified reassignment threshold, *r*, we then retained that vector in the set *SN*. For instance, for *r* = 50%, if more than 50% of *k*-nearest neighbors of the analyzed parameter set belonged to the set *SN*, that parameter set remained in the set *SN*. Otherwise, we re-assigned that parameter vector to the set *SP*. With the proposed reassignment procedure, we emphasized the regions of the parameter space that have a higher proportion of parameter vectors belonging to the set *SN*.

The reassignment procedure introduced two new parameters: the reassignment threshold, *r*, and the number of nearest neighbors, *k*. The values of *r* were chosen on the basis of the initial, unbiased, sampling that was performed in Step III. Specifically, from the initial sampling we could assess the average percentage of *SN* parameter vectors in the set of all vectors. We then set *r* to be a larger than the average percentage so that the parameter classification algorithm could identify the regions in the parameter space with the above than average proportion of *SN* vectors. Assuming that the parameter space was sampled uniformly, we use the parameter *k* to choose the larger or smaller part of the parameter space around the analyzed parameter vector for a possible reassignment. Very large values of *k* are not recommended as the reassignment procedure would consider the overall parameter space and no samples would be retained in the set *SN* as *r* is chosen to be larger than the average percent of *SN* vectors in the overall set of parameter vectors.

### Bayesian inference and parameter classification

Bayesian inference relies on use of Bayes theorem to compute the conditional distribution of a parameter vector *θ* given observed data *x*:

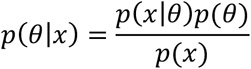

where *p*(*θ*|*x*) is the posterior distribution of the parameters *θ, p*(*θ*) is the prior distribution of parameters, *p*(*x*|*θ*) is the likelihood, and *p*(*x*) is the evidence. In computing the posterior distribution *p*(*θ*|*x*), the evidence can be ignored as it represents a normalizing constant. It is often computationally prohibitive to explicitly evaluate the likelihood function and Approximate Bayesian Computation (ABC) methods are used for approximating this function by simulations (88).

For this type of studies, the ABC rejection algorithm (89) can be used as follows. First, the prior distribution of kinetic parameters is generated using the ORACLE framework or any other method that uses Monte Carlo sampling of uncertain parameters for constructing populations of kinetic models (3–5, 8, 9, 11, 20–28). The corresponding control coefficients are next computed, and the parameter classification algorithm is then used to discard parameter vectors from the prior that gave rise to ambiguous control over analyzed quantities. As a result, the retained samples are distributed according to the approximate posterior distribution of kinetic parameters that give rise to well-determined control over analyzed quantities.

### Computational requirements

The simulations in this study were performed in MATLAB using an Apple MacPro Workstation with 2.7 GHz 12-Core Intel Xeon E5 processor and 64 GB of RAM memory. The required time to generate a set 200’000 kinetic models was ~12.5h, whereas one run of the parameter classification algorithm required several minutes.

## Abbreviations

iSCHRUNK: <u>i</u>n <u>S</u>ilico Approach to <u>Ch</u>aracterization and <u>R</u>eduction of <u>U</u>ncertainty in the <u>K</u>inetic Models
ORACLE: <u>O</u>ptimization and <u>R</u>isk <u>A</u>nalysis of <u>C</u>omplex <u>L</u>iving <u>E</u>ntities
TFA: <u>T</u>hermodynamics-based <u>F</u>lux <u>B</u>alance <u>A</u>nalysis
GEM: <u>GE</u>nome-scale <u>M</u>odel
MCA: <u>M</u>etabolic <u>C</u>ontrol <u>A</u>nalysis

## Acknowledgement

We would like to thank Joana Pinto Vieira for her help with editing this manuscript.

## Supporting information

**S1 File. List of abbreviations for model’s enzymes and chemical species**

**S2 File. Parameter classification rules together with the corresponding feasibility indices.** The rules were obtained from: (i) the parameter classification with the whole set of 258 parameters (sheet “Preliminary training”); (ii) the parameter classification with the preselected 81 parameters (sheet “Training with Fisher preselection”); and (iii) the parameter classification with the top 3 ranked parameters (sheet “Second training (top 3 params)”); (iv) the first parameter classification for the robust ranges of parameters ensuring a negative control for the reference (sheet “Robust Reference”), extreme 1 (sheet “Robust Extreme 1”), and extreme 2 (sheet “Robust Extreme 2”) metabolite concentrations; (v) the second parameter classification with top 3 parameters for the robust ranges of parameters ensuring a negative control for the reference (sheet “Robust Reference Top 3”), extreme 1 (sheet “Robust Extreme 1 Top 3”), and extreme 2 (sheet “Robust Extreme 2 Top 3”) metabolite concentrations; (vi) the parameter classification with reassignment for the reference metabolite concentration ensuring a negative control with the reassignment threshold *r* = 50% (sheet “Reassign – Ref −50%”), and *r* = 30% (sheet “Reassign – Ref −30%”), and ensuring a positive control with *r* = 60% (sheet “Pos Reassign - Ref - 60%”).

**S3 Fig. Preselection of the parameters based on Fisher’s linear discriminant score.** The rules from a tree training with all parameters (blue crosses), and the rules from a tree training with a reduced set of parameters (red diamonds) coincide in the majority of instances.

**S4 Fig. Evolution of PI (blue) together with that of the number of enclosed parameter vectors (orange) for a progressive increase of the lower bound**, 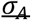, **of the top 9 parameters, while their upper bound was fixed at 1.** For a value of the lower bound 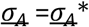, PI was calculated over the range 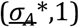 of the parameter *σ_A_*. For example, for 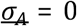, the whole range, i.e., *σ_A_* ∈ (0,1), of a parameter was considered, and the corresponding PI was calculated. For 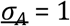, PI was calculated for a fixed value *σ_A_* = 1.

**S5 Fig. Top ranked parameters affecting control of hexokinase (HXK) over xylose uptake (XTR) over three concentrations.** Evolution of the ranking score for the top 10 parameters as a function of the considered number of rules.

**S6 Fig. Metabolic pathways of the VTT C-10880 *S. cerevisiae* strain.** The network includes 102 reactions and 96 metabolite concentrations distributed over cytosol, mitochondria, and extracellular environment. VTT C-10880 strain can consume xylose through the integrated xylose reductase/ xylitol dehydrogenase pathway.

**S7 File. Reference metabolite flux vector together with the metabolite Reference, Extreme1, and Extreme2 concentration vectors.**

**S8 File. Stoichiometry of used models.** List of reactions and the corresponding mass balances.

**S9 File. List of reactions together with the used kinetic mechanisms** (together with S10 File).

**S10 File. Rate expressions for used kinetic mechanisms together with the expressions for the corresponding metabolite elasticities.**

**S11 Fig. Parameter vectors that correspond to a well-determined control for the three metabolite concentration vectors (Reference, Extreme1 and Extreme2).** Parameter sets corresponding to a well-determined control for the three metabolite concentrations (red stars), for two out of three (yellow stars) and for one out of three (blue stars) metabolite concentrations. The gray stars denote the parameter vectors not belonging to a top rule for any of the three concentrations.

## Notes

#### Summary of Updates

The title and the abstract have been modified. The discussion section has been introduced.

